# Comparative Transcriptomic Analysis Revealed Complex Molecular Mechanisms Underlying Pests, Pathogens Resistance and Seed Development in Wild and Cultivated Blackgram

**DOI:** 10.1101/2020.11.09.374041

**Authors:** Avi Raizada, Souframanien Jegadeesan

## Abstract

Blackgram is a widely cultivated pulse crop in Asia. Bruchid pests and yellow mosaic disease (YMD) causes huge loss in legume production including blackgram. Blackgram wild accession (*Vigna m*ungo var. *silvestris*), Trombay wild urd (INGR10133) conferred resistance to bruchids especially *Callosobruchus maculatus*, through antibiosis. However, the mechanisms still remains uncharacterized. We performed the comparative transcriptome analysis of the developing seeds of wild and cultivated blackgram with contrasting phenotypes for 3 traits, bruchids infestation, YMD and seed size. In this study,715differentially expressed genes(DEGs) were re-annotated with reference to NCBI nr database. RNA-Seq was validated by quantitative real-time PCR for 22 DEGs. In Trombay wild, defense related components such as acid phosphatase, vicilins, trypsin inhibitor, brassinosteroid signalling components were found up-regulated. While in cultivar, transcripts for *LEA*, cysteine protease, autophagy related proteins(*ATG3, ATG5, ATG8C and ATG1t*), *DnaJ*, tobamovirus multiplication protein, downy mildew resistance protein, LRR/F-box proteins were found up-regulated. In TW, three transcripts were found common for both bruchids pest and geminivirus resistance (LRR receptor kinase, transmembrane protein 87b and thaumatin like protein).Our study is the first report on transcriptomic differences between wild and cultivated blackgram with new insights into the molecular networks underlying seed development, resistance to pests and pathogens.

## Introduction

Blackgram [*Vigna mungo* (L) Hepper] is an important pulse crop domesticated from *Vigna mungo* var. *silvestris*. It is extensively cultivated for its proteinaceous seeds and as a component of various cropping systems in South Asian regions including India, Myanmar, Pakistan, Bangladesh and Thailand and highly demanded by the sprout industry of Thailand and Japan [1].Yield potential of grain legumes depends on seed features and tolerance to biotic and abiotic stresses. Production of blackgram is severely affected by pathogens in field and by pests during storage especially by Yellow mosaic disease (YMD) [2] and bruchid pests [3] respectively. YMD is the most devastating disease of legumes caused by *Yellow mosaic virus* (YMV) and spread by vector whitefly (*Bemisia tabaci*) while among bruchid pests, *Callosobruchus maculatus* are the most common damage causing agents of stored seeds. In blackgram, among several reported YMD resistance sources, TU94-2 is a well-known elite cultivar resistant to MYMV (*Mungbean yellow mosaic virus*) [4] and among very few bruchid resistance sources reported [5,6], wild accession(*V. mungo* var. *silvestris*)is a well-known resistance source[7] but remained unstudied. Trombay wild blackgram (INGR10133) (TW) is one of the blackgram wild accessions studied in this report which is native to Trombay hills, India. In TW, the resistance trait is controlled by two dominant duplicate genes and resistance mechanism was observed to be larval antibiosis with the constitutive expression of resistance factors resulting in reduced survival, longer developmental period (88 days as compared with 34 days on TU 94-2) and reduced body weight of *C. maculatus* [8,9].Compared to other legumes, very few reports are available in blackgram related to YMD and bruchids[4,5,8,10-14]. Very few studies attempted to understand the transcriptome dynamics of blackgram upon YMV and bruchids attack [15-19] resulting in identification of several defense genes such as defensin, pathogenesis related protein (*PR*) and lipoxygenase (*LOX*) that might be involved in resistance mechanism to pests and pathogens. Through these studies, a foundation to future research has been laid but the molecular mechanisms involved in the resistance to YMV and bruchids still remain incompletely understood. Next-generation sequencing (NGS) technology has been widely used in plant biology for understanding of plant responses under various conditions and in different genetic background [19,20]. Advances in sequencing technology especially RNA-Seq have presented opportunities for comparative transcriptome profiling [17,21,22] for both model and non-model organisms. Several studies reported plant responses to oviposition/bruchids, which are under inducible expression such as neoplastic tissue growth to cast off eggs [23], synthesis of ovicidal compounds [24], release of volatile substance from the leaves that attract parasitoids to kill the eggs [25] and activation of defense genes in response to elicitors such as oral/ovipositor secretions which acts as herbivore associated molecular patterns (HAMPs) [17,26] but no reports are available on constitutive expression of defense response genes in absence of bruchids.

Plant employed growth-defense trade-off scheme for appropriate use of limited resources according to prevailing conditions such as pathogens/pests attack which leads to suppressed growth and development and activation of defense responses to cope up [27,28]. Numerous studies claimed the involvement of several hormone pathways and leucine-rich repeat receptor kinases in defense response [28-31], very little is known about molecular mechanisms regulating growth-defense trade-offs. Here, we report the comparative transcriptome analysis of wild and cultivated blackgram with contrasting phenotypes for 3 traits: YMD, bruchid resistance and seed size (TW −2gms/100 seeds and TU94-2 −4.5gms/100seeds). Present study was conducted on developing seeds from YMD and bruchid non-infected plants, which aimed at exploring transcriptional network and genes involved in resistance mechanism under constitutive expression. We also validated the RNA-Seq results through quantitative real-time PCR (qRT-PCR). Moreover, this study enlightens the transcriptional differences related to innate immune system and seed development of wild and cultivated blackgram.

## Materials and Methods

### Plant material

The wild accession of blackgram(*Vigna mungo* var. *silvestris*) Trombay wild and TU94-2 (*Vigna mungo* var. *mungo*) are maintained at Nuclear Agriculture and Biotechnology Division, Bhabha Atomic Research Centre, Trombay, Mumbai, India. They were grown at the experimental field facility of the Institute at Trombay, Mumbai(latitude 18:54N, longitude 72:49E).

### Transcriptome sequencing, assembly, annotation, DEGs prediction and characterization

Sample preparation from 12 immature seeds of 4 different plants of Trombay wild and TU94-2 harvested 4 weeks after flowering and sequencing was done on Illumina MiSeq platform in a single lane using paired end sequencing chemistry (Xcelris Genomics Ltd. Ahmedabad) [14,32]. Sequencing raw data was processed further to remove adaptor contamination and low quality reads (QV<20) using Trimmomatic v0.30 and high quality reads were assembled using CLC Genomics Workbench with default parameters [14,32]. After assembly, sequence data was submitted to NCBI database with ID. SRR 5931432, SRX3091690 (study accession SRP115376) for TW and SRR 1616991, SRX710526 (study accession SRP 047502) for TU94-2 by the same authors [14,32]. CDS were identified using ORF-predictor with the selection of longest frame and annotated using BLASTx with reference to green plant database with significant similarity (evalue ≤1e-5) [14,32]. After annotation CDS were mapped to Gene Ontology (GO) and KEGG (Kyoto Encyclopedia of Genes and Genomes) database (http://www.genome.jp/kegg). DEGs identification using CLC and FPKM calculation and classification as up and down regulated based on log fold change (FC) values were also done and reported by the same authors [14,32].

### Reannotation of DEGs

In the previous study, on wild blackgram transcriptome sequencing, CDS (DEGs) identified through comparison between wild and cultivated blackgram were annotated based on green plant database which resulted in assignment of majority of DEGs as uncharacterized or hypothetical proteins with undefined function [32]. In this study, DEGs ranging from 3 to 12 and −3 to −12 (range selected randomly) were reannotated using NCBI database which led to the assignment of most of the DEGs with defined function which were left uncharacterized in previous study [32]. Further, after reannotation, based on literature survey, DEGs were selected for their role in pests and pathogenesis defense response or seed development.

### Validation of RNA Seq by Quantitative Reverse Transcription Polymerase Chain Reaction (qRT-PCR)

Total RNA was extracted from immature seeds of Trombay wild and TU94-2 plants with the help of Spectrum Plant Total RNA Kit (Sigma-Aldrich, USA) and treated with DNAse-I (Sigma-Aldrich, USA) to eliminate traces of genomic DNA. The cDNA first strand synthesis was done using a PrimeScript RT-PCR Kit (Clontech, USA) and quantitative real time PCR was performed following manufacturer’s instructions given in SYBR1 Premix ExTaq (TliRNAse H Plus) (Clontech,USA). The PCR amplification was carried out in Rotor-Gene-Q Real-Time PCR System (Qiagen, USA) with the following program, 95°C for 5 min followed by 35 cycles of 94°C for 30 sec, 62°C for 20 sec and 72°C for 20 sec followed by melting of PCR products from 65°C to 95°C. Quantitative real-time PCR experiments for all gene-specific primers were performed twice with three biological replicates run in triplicate. The relative gene expression levels were calculated by relative quantification (RQ) through the 2^-ΔΔCt^ method [33].

## Results

### Transcriptome sequencing, assembly, annotation, DEGs prediction and reannotation

Results of sequencing, assembly, submission of Seq data, annotation and DEGs prediction for TU94-2 and TW developing seeds transcriptomes were reported by the same authors in previous publications[14,32]. DEGs that lacks annotation in previous study (Green plant database) were reannotated with reference to NCBI nr database in this study and assigned with functions (S1Table). A total of 682 DEGs were reannotated including 264 up-regulated (3 to 12 fold change) and 418 down-regulated (−3 to −12 foldchange). Heat map showed the differential expression pattern of top 100 DEGs in Fig. 1.

**Figure 1.**
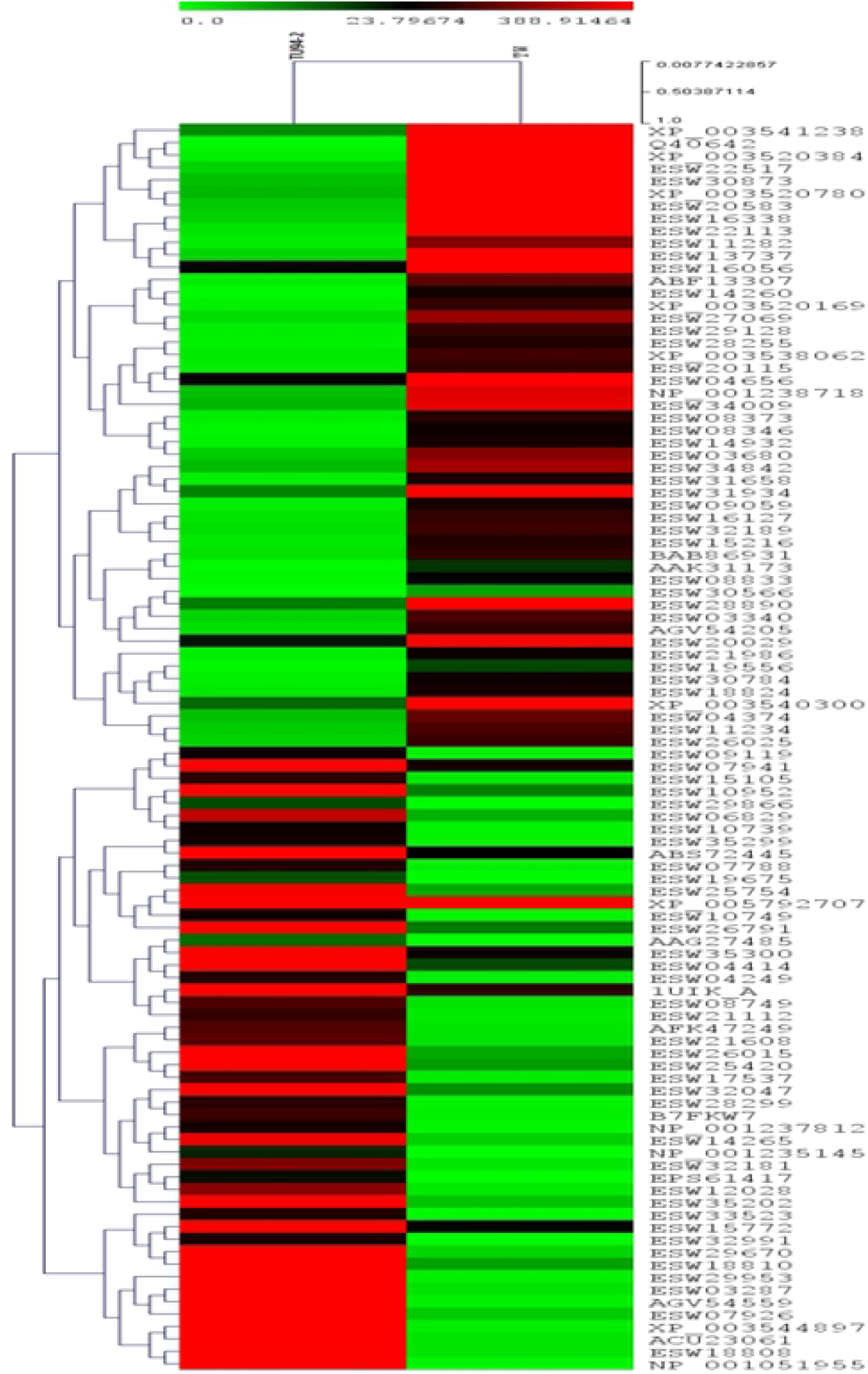
Heat Map showing differential expression pattern of top 100 coding sequences (CDS) of TU94-2 cultivar and Trombay wild (TW) blackgram. Green colour denotes down-regulation and red colour denotes up-regulation of CDS.

### Transcriptome characterization of wild blackgram (Trombay wild)

The transcriptome analysis showed up-regulation of genes associated with pathogen/elicitors perception, downstream signalling and effectors. DEGs related to bruchid pest resistance that were found up-regulated in TW developing seeds are described in Table 1 and DEGs involved in defense responses to other phytopathogens are given in S2 Table.

**Table 1.**
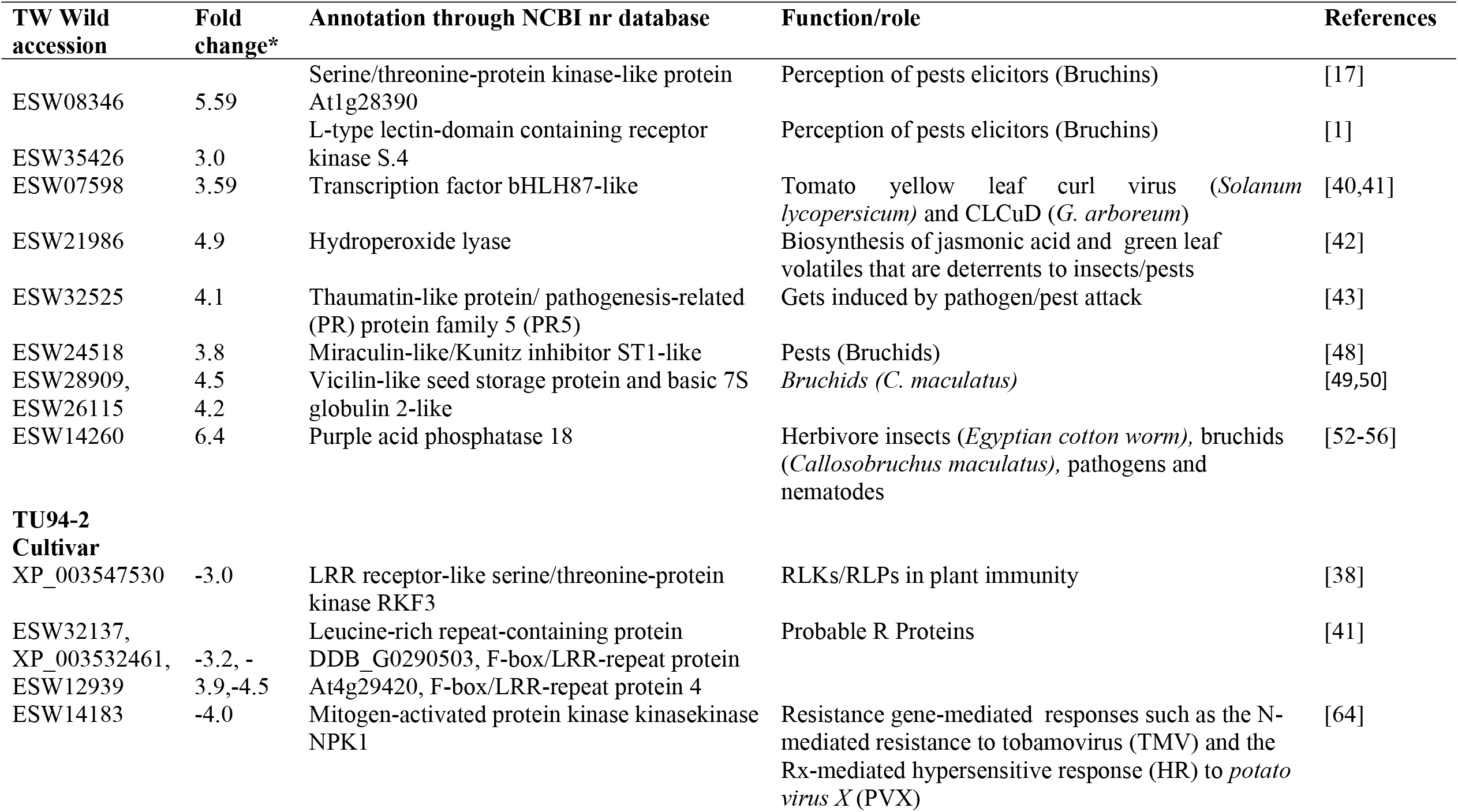
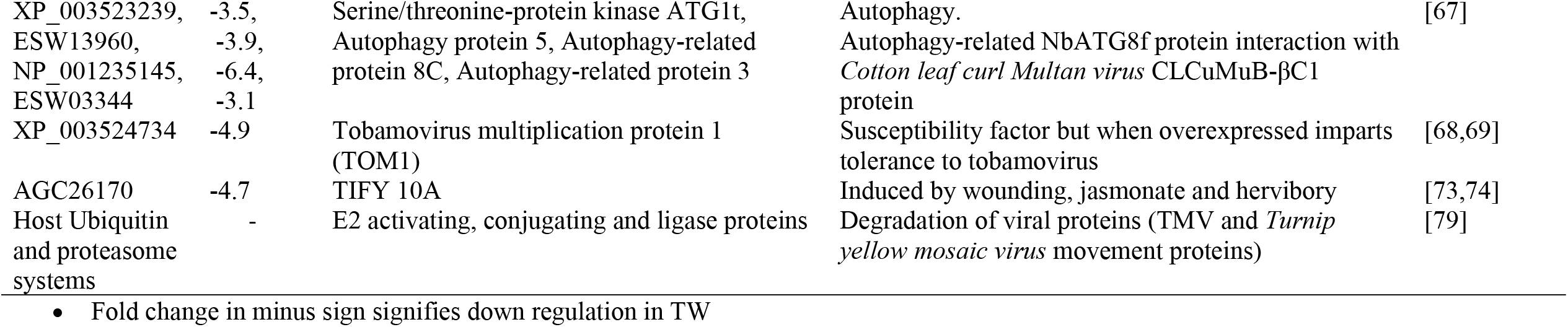
DEGs found (RNA-Seq) to be upregulated in developing seeds of TWwild accession and TU94-2 cultivar involved in pests (Bruchids) and pathogens (Geminiviruses) defense responses.

| TW Wild accession | Fold change* | Annotation through NCBI nr database | Function/role | References |
| --- | --- | --- | --- | --- |
| ESW08346 | 5.59 | Serine/threonine-protein kinase-like protein At1g28390 | Perception of pests elicitors (Bruchins) | [17] |
| ESW35426 | 3.0 | L-type lectin-domain containing receptor kinase S.4 | Perception of pests elicitors (Bruchins) | [1] |
| ESW07598 | 3.59 | Transcription factor bHLH87-like | Tomato yellow leaf curl virus (*Solanum lycopersicum*) and CLCuD (*G. arboreum*) | [40,41] |
| ESW21986 | 4.9 | Hydroperoxide lyase | Biosynthesis of jasmonic acid and green leaf volatiles that are deterrents to insects/pests | [42] |
| ESW32525 | 4.1 | Thaumatin-like protein/ pathogenesis-related (PR) protein family 5 (PR5) | Gets induced by pathogen/pest attack | [43] |
| ESW24518 | 3.8 | Miraculin-like/Kunitz inhibitor ST1-like | Pests (Bruchids) | [48] |
| ESW28909, | 4.5 | Vicilin-like seed storage protein and basic 7S | Bruchids (*C. maculatus*) | [49,50] |
| ESW26115 | 4.2 | globulin 2-like |  |  |
| ESW14260 | 6.4 | Purple acid phosphatase 18 | Herbivore insects (*Egyptian cotton worm*), bruchids (*Callosobruchus maculatus*), pathogens and nematodes | [52-56] |
| **TU94-2 Cultivar** |  |  |  |  |
| XP_003547530 | -3.0 | LRR receptor-like serine/threonine-protein kinase RKF3 | RLKs/RLPs in plant immunity | [38] |
| ESW32137, |  | Leucine-rich repeat-containing protein | Probable R Proteins | [41] |
| XP_003532461, | -3.2, - | DDB_G0290503, F-box/LRR-repeat protein |  |  |
| ESW12939 | 3.9,-4.5 | At4g29420, F-box/LRR-repeat protein 4 |  |  |
| ESW14183 | -4.0 | Mitogen-activated protein kinase kinasekinase NPK1 | Resistance gene-mediated responses such as the N-mediated resistance to tobamovirus (TMV) and the Rx-mediated hypersensitive response (HR) to *potato virus X* (PVX) | [64] |
| XP_003523239, | -3.5, | Serine/threonine-protein kinase ATG1t, | Autophagy. | [67] |
| ESW13960, | -3.9, | Autophagy protein 5, Autophagy-related | Autophagy-related NbATG8f protein interaction with |  |
| NP_001235145, | -6.4, | protein 8C, Autophagy-related protein 3 | *Cotton leaf curl Multan virus* CLCuMuB-βC1 |  |
| ESW03344 | -3.1 |  | protein |  |
| XP_003524734 | -4.9 | Tobamovirus multiplication protein 1 (TOM1) | Susceptibility factor but when overexpressed imparts tolerance to tobamovirus | [68,69] |
| AGC26170 | -4.7 | TIFY 10A | Induced by wounding, jasmonate and herbivory | [73,74] |
| Host Ubiquitin and proteasome systems | - | E2 activating, conjugating and ligase proteins | Degradation of viral proteins (TMV and *Turnip yellow mosaic virus* movement proteins) | [79] |

- Fold change in minus sign signifies down regulation in TW

The transcripts of three LRR-RLKs (leucine-rich repeat receptor-like protein kinase); Leucine-rich repeat receptor-like protein kinaseAt1g68400, LRR receptor kinase *BAK1* or somatic embryogenesis receptor kinase 1-like*(SERK)* and one each of LRR receptor-like serine/threonine-protein kinase *ERECTA*, RPK; receptor serine/threonine-protein kinase-like protein At1g28390, L-type lectin-domain containing receptor kinase S and transmembrane protein 87B were over-expressed. Four genes related to signal transduction were found up-regulated, of which 2 were serine/threonine-protein kinases:Serine/threonine-protein kinase *STY8* isoform C, probable serine/threonine-protein kinase At4g35230 and 2 were calcium related proteins: Calmodulin-like protein 1(*CML1*) and calnexin homolog isoform X1. Genes encoding several ribosomal proteins, translation initiation factors, carbohydrate metabolism genes, tubulin chains and myosin binding proteins, cell wall proteins such as glycosyltransferases, arabinogalactans and expansins were highly expressed. Stress-associated genes such as heat shock proteins, stress response proteins, DnaJ homologs, chaperons and defense-associated genes such as acid phosphatase*(ACP)*, 7S globulins, vicilins, thaumatin, miraculin and thioredoxin which are reported effectors against pathogens and pests were up-regulated. Transcripts of abscisic acid receptor*(PYL9)*, protein phosphatase 2C (*PP2C*) and abscisic acid insensitive 5 (*ABI5*)were enriched. Interestingly gene encoding hydroperoxidelyase (*HPL, CYP74B*) which is a cytochrome P450 present in chloroplasts was found to be upregulated. Several transcriptional regulators belonging to different families were found up-regulated, which included ethylene-responsive transcription factor *ERF061*, zinc finger A20 and AN1 domain-containing stress-associated protein 5, transcription factor *LHW*, transcription termination factor *MTEF18*, RING-H2 finger protein *ATL8* and DNA-directed RNA polymerase III subunit *RPC7*. Several DEGs involved in growth, development, carbohydrate and amino-acid metabolism were found up-regulated. DEGs related to seed development included *GID1* receptor, *DELLA* proteins and Caffeic acid 3-O-methyltransferase were found up-regulated.

### Transcriptome characterization of cultivated blackgram(TU94-2)

Differentially expressed genes involved in defense responses to geminivirus and found up-regulated in TU94-2 developing seeds are described in Table 1 and DEGs involved in defense responses to other phytopathogens are given in S3 Table. Transcriptome analysis showed basal level up-regulation of genes encoding pathogen recognition receptors (PRRs) and LRR-containing proteins such as LRR receptor-likeserine/threonine-protein kinase *RKF3*, receptor-like protein kinase *FERONIA*, LRR containing protein DDB_G0290503, two F-box/LRR-repeat proteins(protein At4g29420and protein 4) and four uncharacterized transmembrane proteins(transmembrane protein 205, transmembrane 9 superfamily member 2, transmembrane protein 230 and transmembrane protein *PM19L*) (S3 Table). The transcripts of upstream regulator of *TOR*, 1-phosphatidylinositol-3-phosphate 5-kinase *FAB1B*, several downstream effectors of TOR including abscisic acid receptor *PYL12*, protein phosphatase 2C (*PP2C*), three autophagy related proteins (*ATG 3,5* and *8C)* and several serine/threonine proteine kinases, which constitutes TOR signaling such as Serine/threonine-protein kinase *AtPK2/AtPK19, ATG1t, CTR1, PBL11,PAKD*and *HT1* were upregulated. Interestingly gene encoding chloroplast located lipoxygenase (*LOX*) which is involved in biosynthesis of jasmonic acid was up-regulated. Noteworthy transcripts encoding components of PTI and ETI signaling including calcium-dependent protein kinase 28 (*CPK28*), mitogen-activated protein kinase kinase 5, mitogen-activated protein kinase kinase kinase *NPK1*, tobamovirus multiplication protein 1 (*TOM1*) were found to be over-expressed. A number of transcription factors associated with pathogenesis were also found upregulated which belonged to families such as *MYB, NAC* and *WD*-repeat domains containing transcription factors. Several transcripts of *DNAJ*, heat shock chaperones, ubiquitin activating, conjugating enzymes, ligases, SEC interacting proteins and proteasomes were observed to be highly expressed. In addition, another susceptibility factor found to be enriched in TU94-2 cultivar was DOWNY MILDEW RESISTANCE 6(*DMR6*). Similar to TW, DEGs involved in growth, development and metabolism were also found up-regulated in TU94-2 developing seeds. Apart from this DEGs related to cell expansion, cell wall loosening and protein turnover (*MYB1R1* DNA-binding protein, xyloglucan endotransglucosylase, Ubiquitin proteasome system components and rhamnogalacturonan I rhamnosyltransferase 1) which directly or indirectly regulates seed development were also found up-regulated in TU94-2.

### Validation of differentially expressed genes using quantitative real-time PCR

We quantified relative expression of total 27 genes, 22 DEGs from the RNA-seq dataset of blackgram developing seeds and 5 genes coding for putative factors reported for imparting resistance to pests were based on literature survey. The elongation factor EF 1α gene was used as an internal control for normalisation. Details of primers used in this study are given in Table 2.The relative gene expression patterns of the qRT-PCR results for 22 genes were consistent with RNA-Seq data.Out of 22 genes, 15 up-regulated and 7 down-regulated genes in TW were validated by qRT-PCR (Fig. 2(a,b)). TW transcript coding for an acid phosphatase protein (Purple acid phosphatase *ACP18*) had higher (1.3 fold) expression levels compared to TU-94-2 cultivar (Fig. 3a). The transcript of universal stress protein *PHOS32* showed 2.5 fold more expression in wild than cultivar under controlled conditions (Fig. 3b). We also validated a leucine-rich repeat receptor like kinase gene (*LRR-RLK*) which showed enhanced expression (1.4 fold change) in wild in comparison to cultivar. Similar fold changes were also observed in TW as compared to TU94-2 cultivar for the following under controlled conditions, which included golgin subfamily A member 6-like protein 6, multiple organellar RNA editing factor 2, RING-H2 finger protein ATL8-like, ANTAGONIST OF LIKE HETEROCHROMATIN PROTEIN 1, protein RETICULATA-RELATED 1 (chloroplastic), EID1-like F-box protein 2, prostatic spermine-binding protein, protein *PNS1*, glycosyltransferase *BC10*, gibberellin receptor *GID1*, geranylgeranyl pyrophosphate synthase 7 (chloroplastic) and 50S ribosomal protein 5 (chloroplastic) (Fig. 3a and 3b).

**Table 2.**
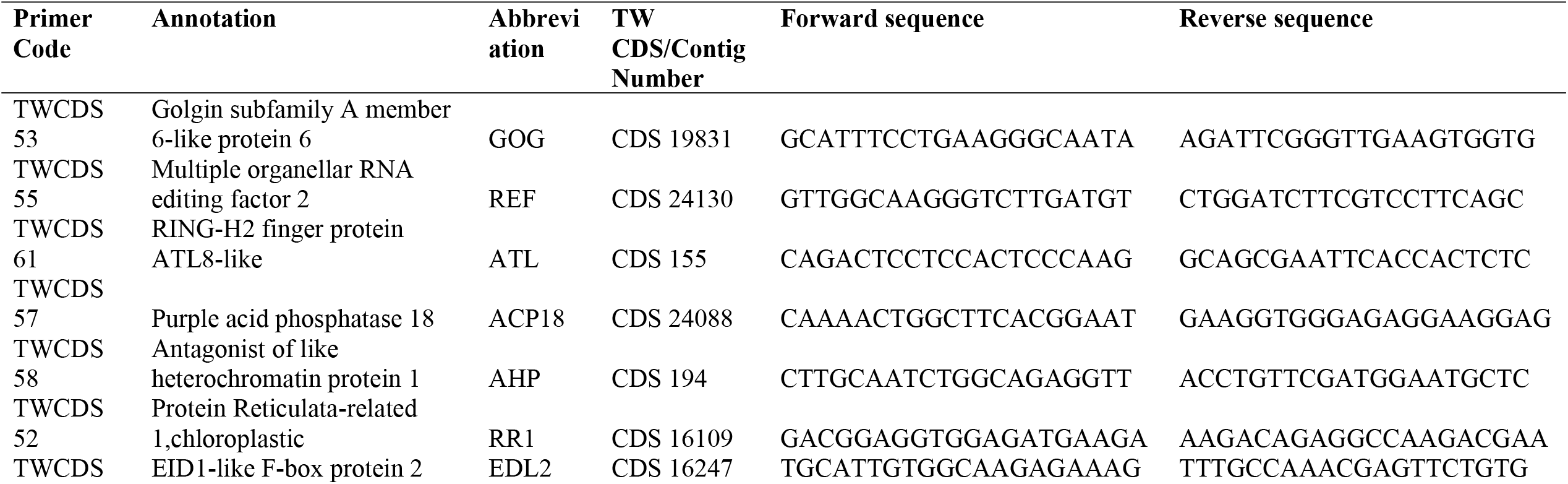
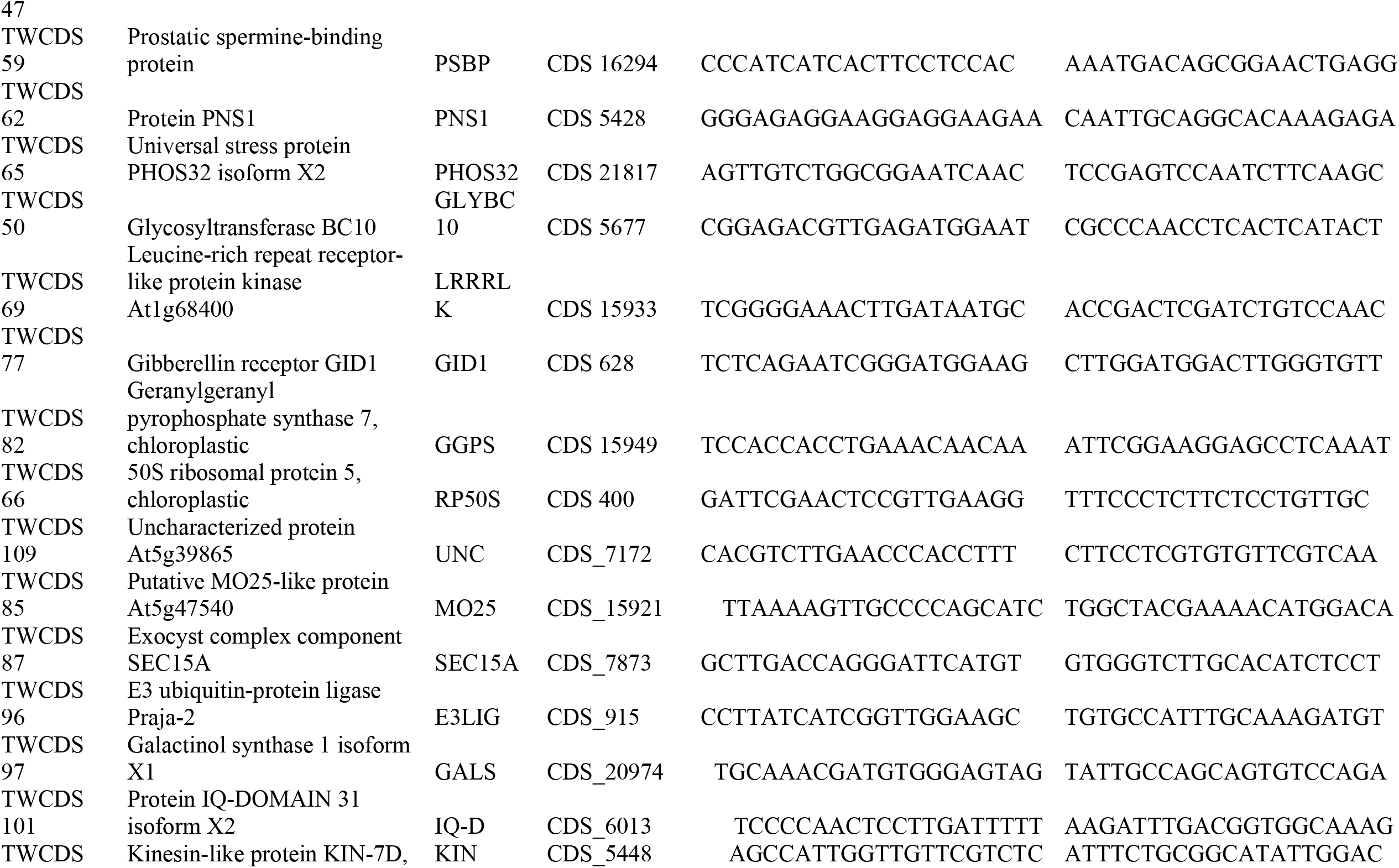
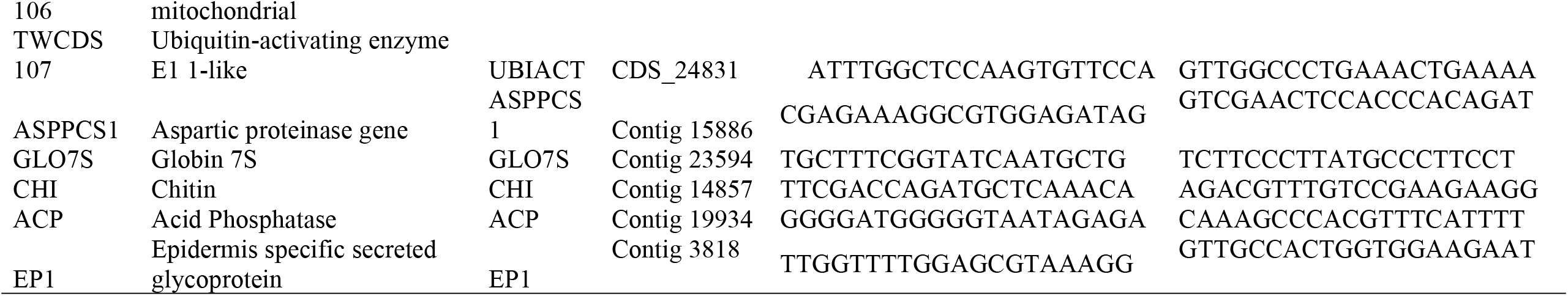
Details of Primers used in quantitative real-time PCR experiment of this study

**Figure 2.**
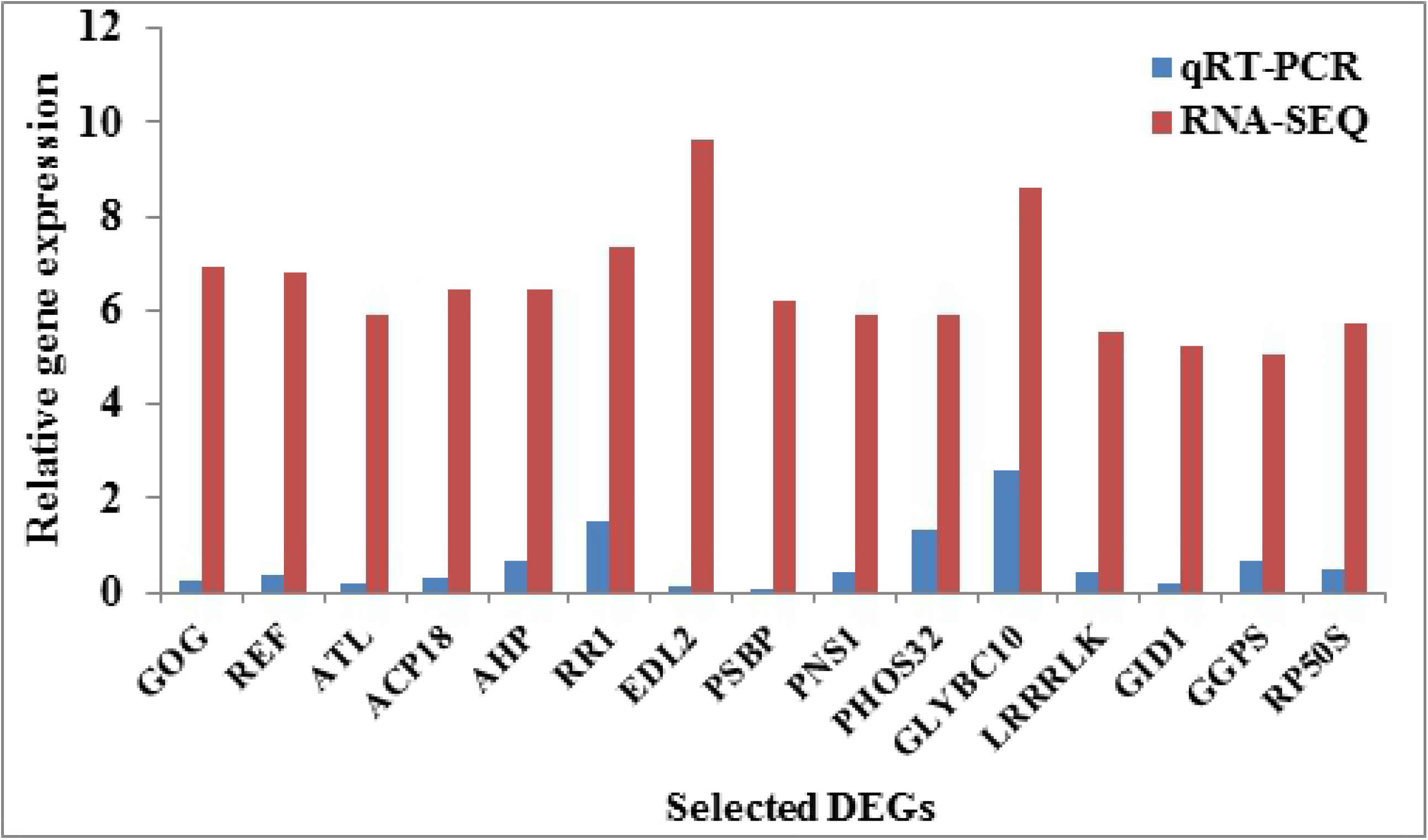
Validation of RNA-Seq result with RT PCR. Expression of 22 randomly selected genes was examined by RT-PCR analysis. a) Expression pattern of 15 up-regulated genes and b) Expression pattern of 7 down-regulated genes. For each gene, fold changes were calculated by ^ΔΔ^Ct method in the RT-PCR, converted to log values and with the FPKM values (log FPKM TW/TU94-2) in the RNA-Seq.

**Figure 3.**
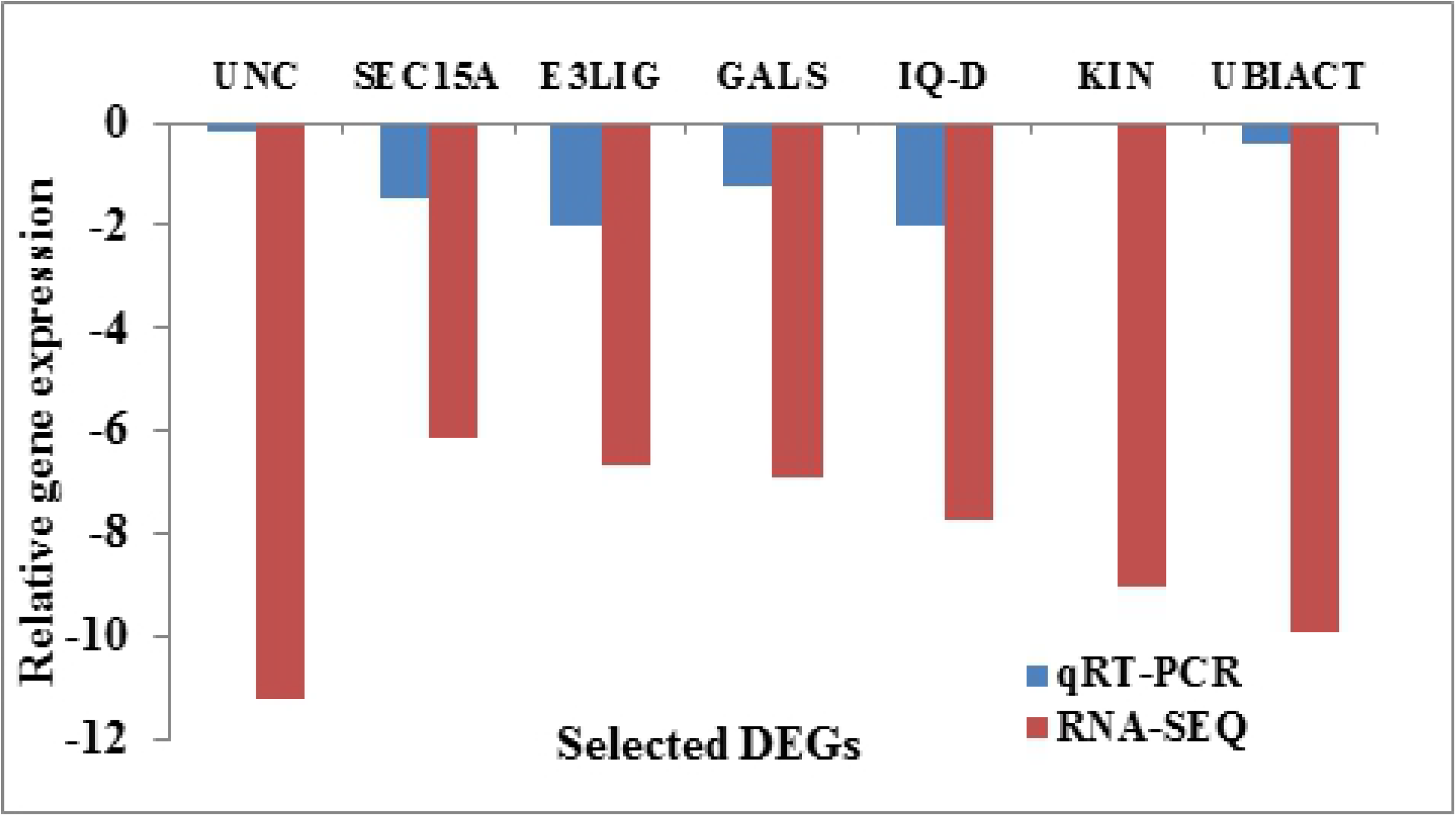
Expression levels of twenty genes found up-regulated in TW non-infested developing seeds. The expression levels were normalized with the help of EF1*α* gene of blackgram. Expression levels were calculated by ^ΔΔ^Ct method in the RT-PCR but not converted to log values. a) and b) showed Fold changes obtained only from qPCR experiments and calculated through ^ΔΔ^Ct method. Full names of genes abbreviations were given in primer details Table 2. Bars represent mean ± standard deviation.

### Comparative analysis of TW and TU94-2 transcriptomes with respect to bruchid, geminiviruses resistance and seed development

In TW seed transcriptome, 11 up-regulated transcripts encode for cellular factors related to resistance against bruchid pests and 5 up-regulated transcripts encode for geminiviruses resistance related factors. Among these, 3 transcripts (LRR receptor kinase, transmembrane protein 87b and thaumatin like protein) were found to be common which are involved in resistance to both pests and geminiviruses (Fig. 4 and S4Table). Similarly, TU94-2 seed transcriptome showed 15 up-regulated transcripts related to geminiviruses resistance, 9 transcripts for pests resistance and 5 transcripts (LRR receptor-like serine/threonine-protein kinase *RKF3*, leucine-rich repeat-containing protein *DDB_G0290503*, seed linoleate 9S-lipoxygenase-3, cysteine protease and cysteine protease *RD19D*) as common for both agents (Fig. 4 and S4Table).While for seed development, majority of DEGs related to growth and metabolism showed similar pattern of expression. But few DEGs showed considerable difference in the expression level which may be responsible for genotypic specific traits. In TW transcriptome 12 transcripts were found to be up-regulated, which are related to protein synthesis and ubiquitin proteasome machinery and one each for less common genes such as caffeic acid 3-O-methyltransferase, vegetative cell wall protein gp1-like and *DELLA* protein *GAI* (S5Table). Whereas, in TU94-2 transcriptome, 24 up-regulated transcripts code for protein synthesis and ubiquitin proteasome machinery components and one each for rhamnogalacturonan I rhamnosyltransferase 1-like, xyloglucan endotransglucosylase/hydrolase 2, cellulose synthase, galactinol synthase, two for galactosyltransferase and four transcripts for late embryogenesis abundant protein (S5Table).

**Figure 4.**
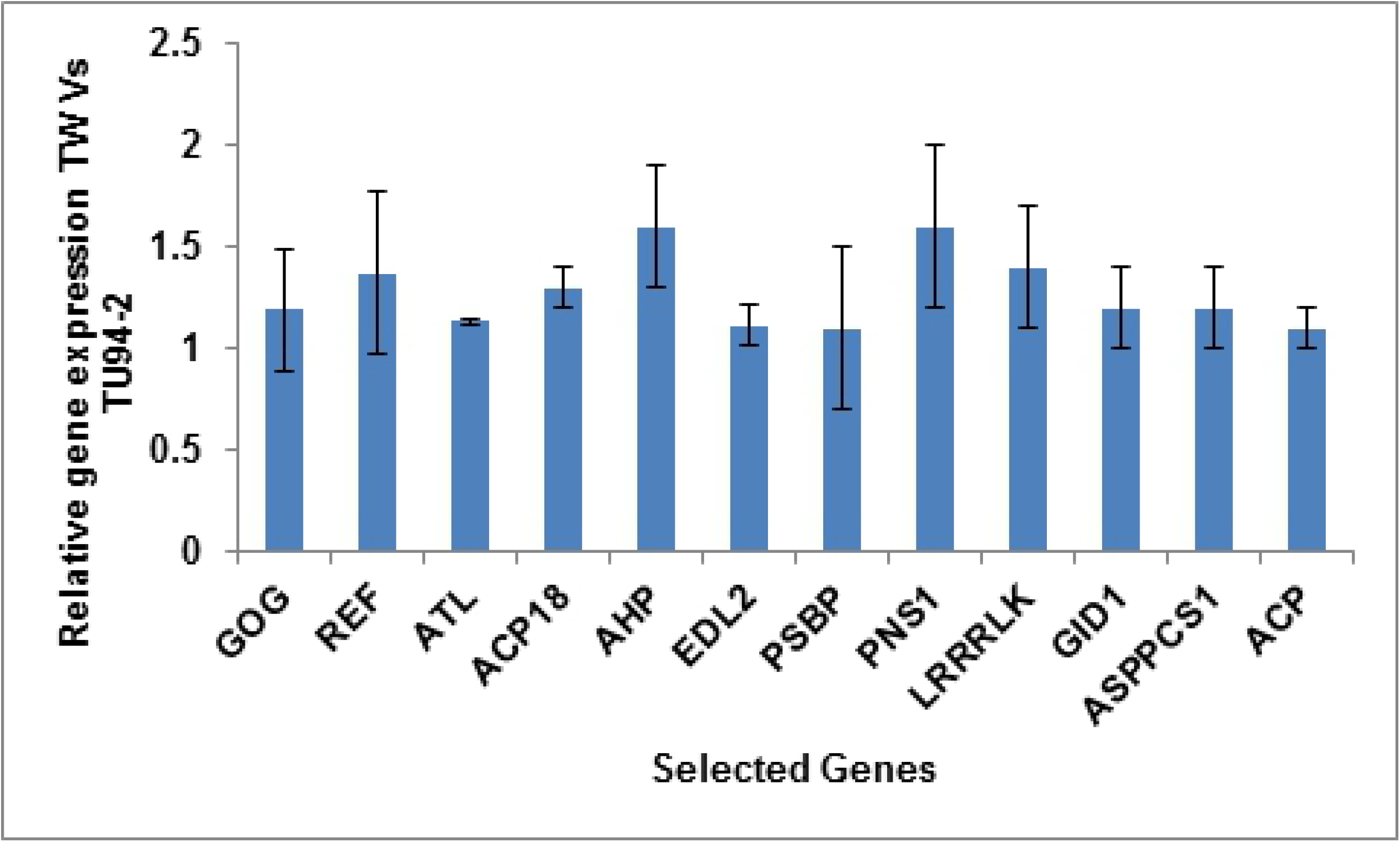
Venn diagram showing up-regulated DEGs of TW and TU94-2 blackgram genotypes related to bruchids pest and geminivirus interaction.

## Discussion

In this study, we compared the transcriptomes of developing seeds of wild and cultivated blackgram differing in phenotypes for 3 traits, to identify the resistance mechanism and candidate genes expressing constitutively at basal level.

### Transcriptome characterization of wild blackgram: Innate immune system in response to bruchid pests and other phytopathogens

Timely perception of pathogens and effective defense response by plant host depends on plasmamembrane localized receptors for elicitor recognization and downstream signaling[34]. In TW, transmembrane receptors *BAK1, SERK1, ERECTA* and lectin domain receptors were found up-regulated, which are regulators of PAMP triggered immunity. For example, *BAK1* is required for immunity to diverse RNA viruses [35-37]. Recently, lectin receptor kinase and chitinase were found to be associated with bruchid resistance trait in wild blackgram accession TC2210 through high-density linkage map [1]. Up-regulation of receptor serine/threonine kinase (*RSTK*) in the oviposited developing seeds of moderately tolerant blackgram is speculated to be required for perception of elicitors (bruchins) [17]. Serine/threonine-protein kinase At4g35230/ BR-signaling kinase 1 *(BSK1)*was also up-regulated which mediates signal transduction from receptor kinase *BRI1*and positively regulates brassinosteroid signaling and plant immunity [38]. Therefore, above up-regulated receptor kinases might be important for resistance to bruchids and pathogens in wild blackgram. The most represented transcription factorsin the DEGs included Ethylene response factors (ERFs), followed by the Tri-helix transcription factors. ERFs are regulators of pathogenesis-related genes and ethylene-, salicylic acid-, and jasmonic acid-inducible genes [39]. The bHLH transcription factor was also up-regulated which imparted immunity to viruses such as tomato *yellow leaf curl virus*[40] and in cotton leaf curl disease (CLCuD) stress [41]. Hydroperoxidelyase transcript (*HPL, CYP74B*) was found up-regulated which is involved in biosynthesis of jasmonic acid and green leaf volatiles that are deterrents to insects/pests [42]. RNA-Seq results showed basal level over-expression of anti-insect and anti-pathogenic compounds such as acid phosphatase, thaumatin like proteins, trypsin inhibitor/miraculin, vicilin and 7s globulin. Thaumatin-like proteins are the pathogenesis-related (PR) protein family 5 (PR5) proteins which are known to get induced by pathogen/pest attack [43]. Proteinase inhibitors (PIs) are natural defense proteins generally present in seeds which gets induced by herbivory or wounding [44] and imparts a cumulative protective effect on plants due to their anti-nature for insects, nematodes, viruses, bacteria and fungi pathogens [45-47]. Trypsin is known to be involved in developmental processes of insects such as molting and synthesis of neuropeptides, therefore trypsin inhibitors can affect these critical stages, which lead to growth and developmental retardation of the larvae [48]. Likewise numerous studies have reported detrimental effects of legume vicilins and 7S globulins on insects development especially for *C. maculatus* [49,50]. Besides being involved in phosphate acquisition and utilization [51], acid phosphatases have been also implicated in resistance to herbivore insects (*Egyptian cotton worm)*[52,53], bruchids (*Callosobruchus maculatus)*[54], pathogens and nematodes [55,56]. Above RNA-Seq results suggests that in wild blackgram developing seeds, up-regulated cellular components primarily functions in seed development and secondarily involved in defense processes. Moreover, different regulation of processes might be responsible for differential expression of defense effectors in TW, thus imparting it enhanced tolerance to specific pests/pathogens and different from cultivated blackgram. Pictorial view of hypothesized pathway and processes operating in TW developing seeds that may be controlling resistance against pests and pathogens is given in S1Fig.

### Transcriptome characterization of TU94-2 cultivated blackgram: Defense system in response to viruses and other phytopathogens

Similar to TW, TU94-2 transcriptome showed up-regulation of defense related genes different from TW. RNA-Seq showed up-regulation of FERONIA *(FER)* that serves as a receptor for a unique peptide ligand, RALF1 (Rapid Alkalinization Factor 1) and play role in effector-triggered immunity (ETI) through the RALF1–FER–RIPK signalling module that may intersect with the RIPK–RIN4 (RPM1-induced protein kinase - RPM1-interacting protein 4) pathway [57,58]. Several uncharacterized RLKs and LRR containing proteins were found up-regulated in TU94-2 developing seeds that may be candidate R genes and may be involved in the perception of geminiviruses and other pathogens. For example, it was shown that C4 protein of TYLCV, BCTV and NSP protein of *cabbage leaf curl virus* interacts with BARELY ANY MERISTEM (BAM) and with LRR receptor like kinase [59-61]. Mitogen-activated protein kinase kinase 5 is a component of MAP kinase signalling cascade (MEKK1, MKK4/MKK5 and MPK3/MPK6) which gets activated by bacterial flagellin receptor FLS2 and stimulate hydrogen peroxide generation during hypersensitive response-like cell death [62,63].Transcript of mitogen-activated protein kinase kinase kinase *NPK1*was found over-expressed, which plays a role in the NACK-PQR (NPK1-NQK1/MEK1-NRK1) MAP kinase signaling pathway and controls resistance gene-mediated responses such as the N-mediated resistance to tobamovirus (TMV) and the Rx-mediated hypersensitive response (HR) to *potato virus X* (PVX)[64]. Plants employ both RNA silencing and autophagy as antiviral defense strategies during geminivirus infection for silencing of viral transcripts and degradation of viral virulence factors respectively [65,66]. The TU94-2 transcriptome showed up-regulation of transcripts for three autophagy related proteins *ATG 3,5* and *8C* and one serine/threonine-protein kinase *ATG1t* that may be involved in interaction with geminiviruses as observed for autophagy-related NbATG8f protein with the *cotton leaf curl multan virus* CLCuMuB-βC1 protein [67].The RNA-Seq data showed enriched transcript of tobamovirus multiplication protein 1, which is a susceptibility factor and necessary for intracellular multiplication of tobamovirus [68] but it’s over expression leads to increased accumulation of the membrane-bound forms and decreased accumulation of the soluble forms thus inhibiting tobamovirus multiplication [69].The family of TFs with the most members represented in DEGs included ERFs, followed by the zinc finger CCCH containing protein, *MYB, WRKY, NAC* and *WD*-repeat families, which regulates several jasmonate and ethylene responsive defense genes under pathogen attack [70] as reported in *G. arboreum* defense against CLCuD[41].Transcript for *ERF9*was up-regulated. *ERF9*binds to the GCC-box pathogenesis-related promoter element under stress [71] and negatively regulates defense against necrotrophic fungi[72]. Gene encoding *TIFY10A*was over-expressed, which is a repressor of jasmonate responses and gets induced by wounding, jasmonate and hervibory [73,74]. Geminivirus infection also induces the expression of a DNA-binding protein TIFY4B that acts as a geminiviral resistance factor. The interaction of CabLCV and TGMV TrAPs with TIFY4B inhibits its potential role in cell cycle arrest [75]. Transcripts for β−1,3-glucanase, DnaJ, heat shock chaperones and callose synthase were up-regulated, which might hinders cell to cell movement of viral particles as observed for β−1,3-glucanase interaction with TGB2 protein of Potato Virus X (PVX) [76] and transportation and replication of geminiviruses as observed for DnaJ (HSP 40) [77,78]. In TU94-2 several transcripts components for ubiquitin proteasome system (UPS) were found up-regulated, that are known to target the virus proteins for degradation as defense strategy [79]. For example, SUMO-conjugating enzyme 1 (SCE1) interaction with geminiviral Rep protein [80] and ubiquitin-conjugating (UBC) enzyme(SlUBC3) interaction with *Cotton leaf curl Multan virus* (CLCuMV) βC1 protein [81]. This suggests that ubiquitin mediated proteolysis could be a defense strategy against symptom development. Gene encoding lipoxygenase *(LOX)* was found to be up-regulated that is known to be involved in jasmonic acid (JA) synthesis and is also induced by wounding, hervibory and pathogen invasion. This leads to induction of genes encoding proteinase inhibitors, flavonoid biosynthesis (chalcone synthase and phenylalanine ammonia lyase), sesquiterpenoid biosynthesis (hydroxymethylglutaryl CoA reductase), thionin (antifungal protein), and osmotin (antifungal protein). Interestingly, another gene coding for immunity suppressor found up-regulated was DOWNY MILDEW RESISTANCE 6 encoding a salicylate-5hydroxylase that converts salicylic acid (SA) to 2,3-dihydroxybenzoic acid (2,3-DHBA) [82]. It negatively regulates defense related genes (e.g. *PR-1, PR-2*, and *PR-5*) and is required for susceptibility to the downy mildew pathogen *Hyaloperonospora arabidopsidis, Pseudomonas syringae* pv. tomato DC3000 and oomycete *Phytophthora capsica* [83]. Likewise, in comparison to TW, different regulatory processes may be responsible for differential expression of defense effectors in TU94-2, thus, imparting it enhanced tolerance to diverse range of pathogens and different from wild blackgram. Pictorial view of a hypothesized pathway and processes operating in TU94-2 developing seeds that may be controlling resistance against geminiviruses and pathogens is given in S2Fig.

### Seed Development

In this study, RNA-Seq results revealed differential expression of genes related to cell wall modification, carbohydrate metabolism, and hormone signalling, which were also reported in seed development studies in legumes [84-86]. In TU94-2 developing seeds, genes controlling seed size/weight, seed coat texture etc., were found to be upregulated which included MYB1R1 DNA-binding protein (*R2R3 MYB, MYB56* and a *MYB*-like DNA-binding protein [20, 87]), xyloglucan endotransglucosylase [20, 88], Ubiquitin proteasome system components [20, 80 89] and rhamnogalacturonan I rhamnosyltransferase 1[81 90]. While, in small sized TW seeds, genes found upregulated and observed in seed development included *GID1* receptor, *DELLA* proteins and Caffeic acid 3-O-methyltransferase. DELLA is an inhibitor of plant growth which gets degraded on application of GA3 hormone on dormant seeds, thereby promoting plant germination [91]. Caffeyl alcohols derived lignins were found in the seed coat of castor bean [92] and early expression of lignin related genes in small-seeded castor bean seed coat leads to early lignin deposition thus restricting seed size due to suppressed cell division by rigid secondary cell walls [93]. Seed development studies revealed that seed coats posed physical constraints on embryo and/or endosperm growth by setting an upper limit for seed size [94,95]. Abscisic acid insensitive 5 (*ABI5*) was found to be upregulated in TW developing seeds which negatively regulates seed germination in *Arabidopsis*[96] through mediating cell responses to ABA in seeds and vegetative tissues.

### Comparative analysis of TW and TU94-2 transcriptome with respect to pests, geminiviruses and seed development

In TW developing seeds, up-regulated transcripts coding for bruchidpest resistance related factors are predominant compared to geminiviruses interaction related factors. In contrast in TU94-2 seeds, up-regulated transcripts related to geminiviruses interaction constitute majorly compared to bruchidpest resistance related factors. Lectin domain containing protein, acid phosphatase, vicilin, thaumatin, miraculin/trypsin inhibitor transcripts found up-regulated in TW are known for bruchid pest resistance as discussed earlier, while cysteine protease, endochitinase, wound-induced protein also related to pests resistance were found up-regulated in TU94-2 developing seeds. This suggests that bruchid resistance trait of TW could be due to Lectin domain containing protein, acid phosphatase, vicilin, thaumatin, miraculin/trypsin inhibitor. Both TW and TU94-2 developing seeds showed presence of several forms of DnaJ protein which is known to play a role in geminivirus multiplication and movement, thus suggesting the less significance of DnaJ proteins for geminivirus resistance in TU94-2.Transcripts coding for LRR repeat proteins and autophagy related proteins, which play a role in interaction with geminiviruses were found to be up-regulated in TU94-2, suggesting their significance in imparting YMD resistance trait to TU94-2. In TU94-2 developing seeds, transcripts related to protein biogenesis, turnover and ubiquitin proteasome system were more prominent compared to TW, thus suggesting more metabolically active state in TU94-2. Moreover, up-regulation of several distinct transcripts related to cell wall modification and texture in TU94-2 developing seeds suggests difference in seed coat texture of both blackgram genotypes.

In conclusion, this is the first report that describes the differences in transcriptomes between wild and cultivated blackgram differing in three important traits. Our analysis defined putative resistance mechanism and candidate genes under constitutive expression in blackgram that are related to defense responses to diverse pests and pathogens. This study lays a theoretical foundation for an improved understanding of the molecular mechanisms involved in resistance to bruchid pest, geminiviruses and other pathogens and mechanisms regulating seed development.

## Authors’ contributions

AR carried out the experiment and wrote the manuscript with support from JS. JS conceived the idea and supervised the project. Both authors have read and approved the manuscript.

## Corresponding author

Correspondence to

---

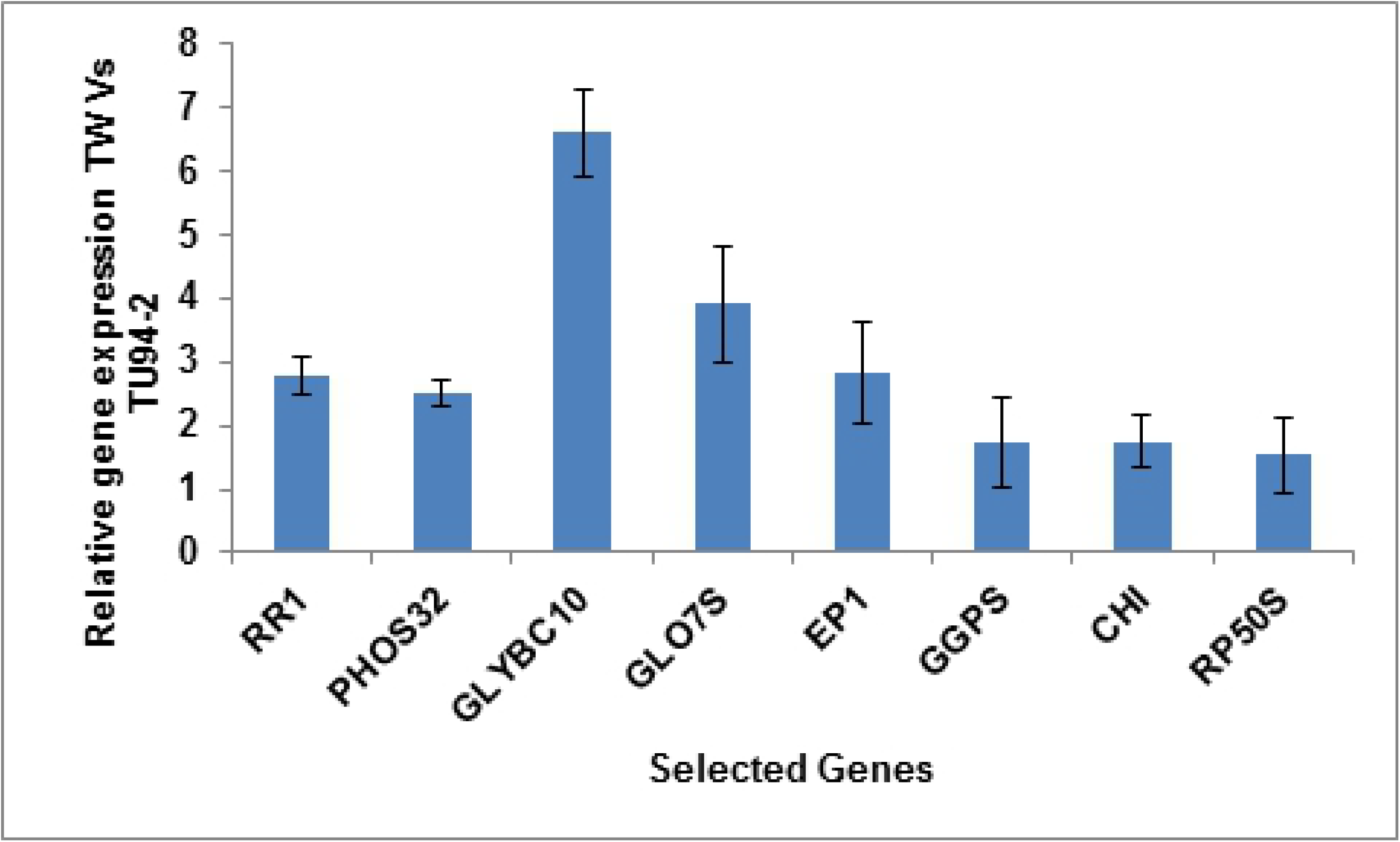

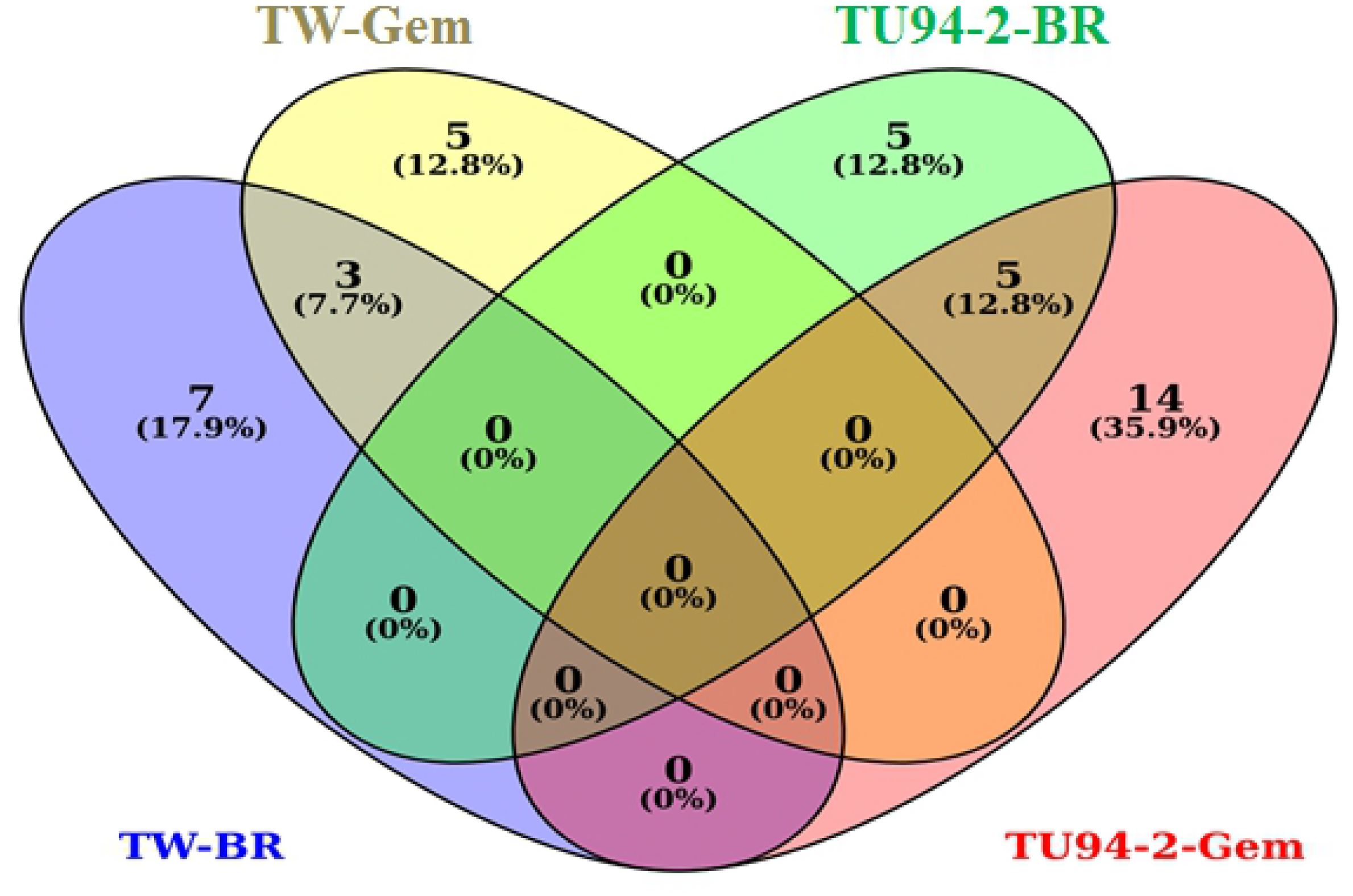

